## Supplementary Figures S1 and S2 for "Dynamics of the viral community on the surface of a French smear-ripened cheese during maturation and persistence across production years"

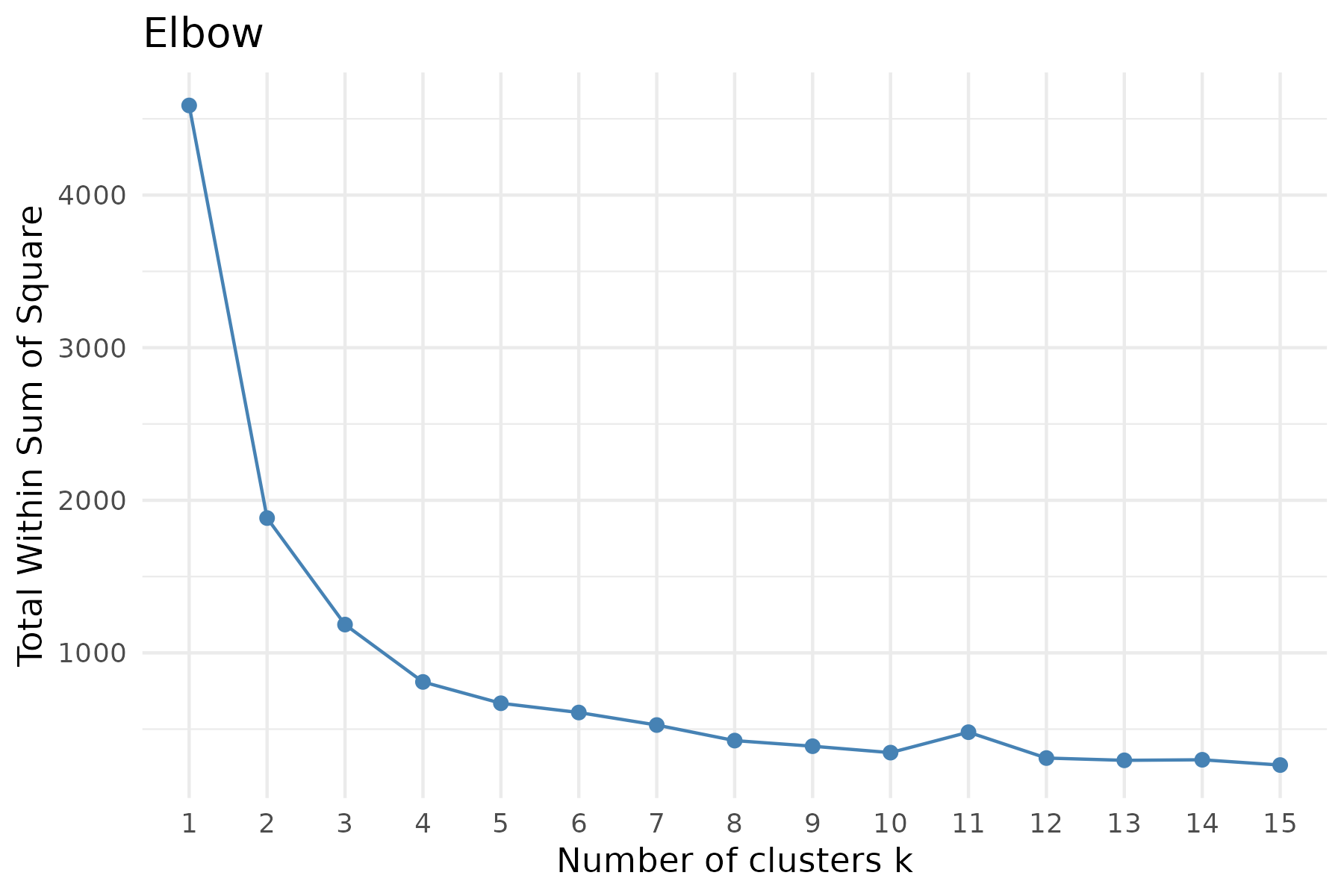

**Figure S1: Estimation of the optimal number of vOTUs clusters in the short-term dataset.** The total within-cluster-sum-of-square was calculated for each number of clusters and the elbow point was used to determine the optimal number of clusters.

**Figure S2: Differential abundance analysis of the virome data between the two stable stages represented by W1-2 and W4-5 samples (short-term study).** Log2 fold change values obtained for differential abundant vOTUs (dots) are represented and the dot colour indicate the predicted host genus.

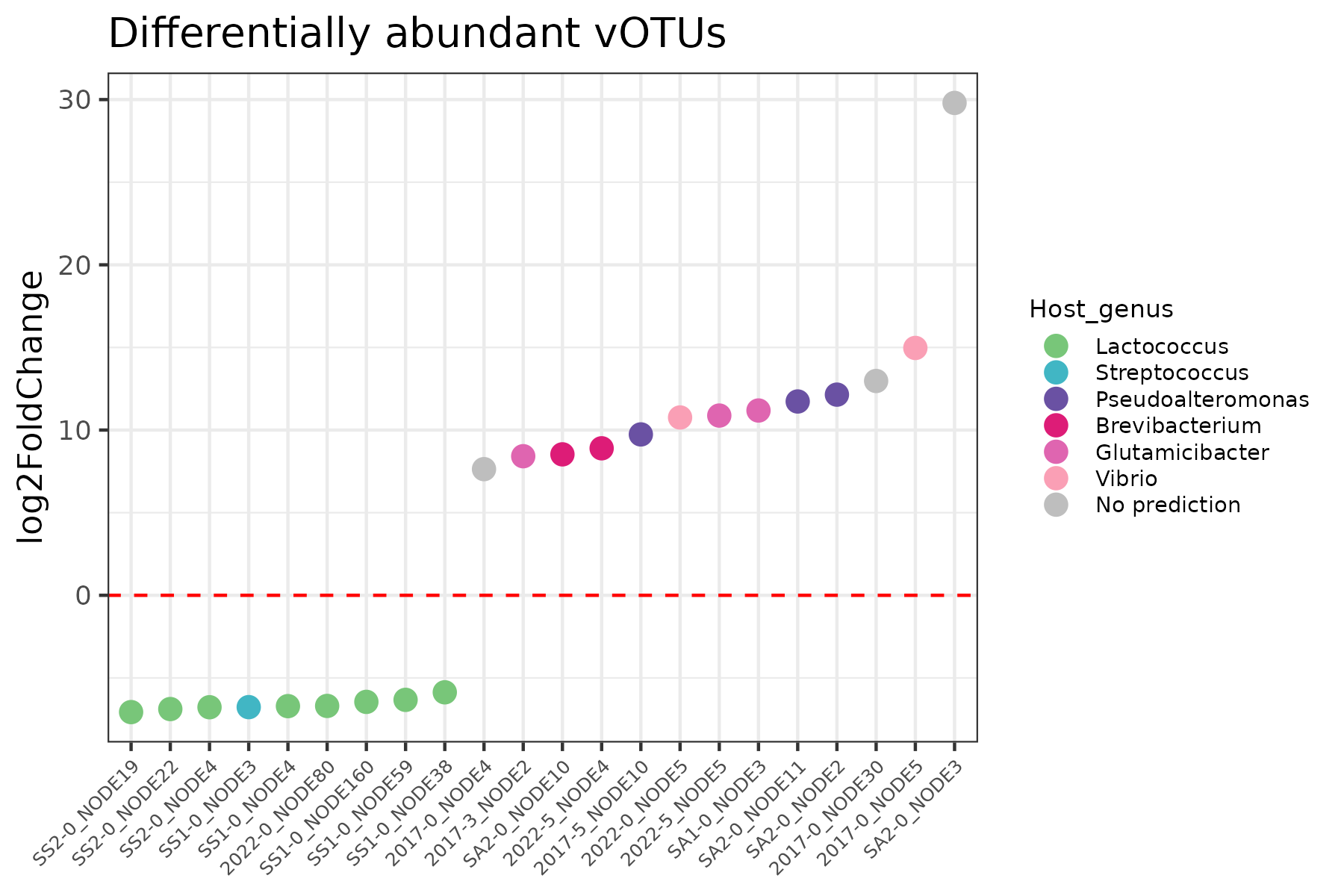

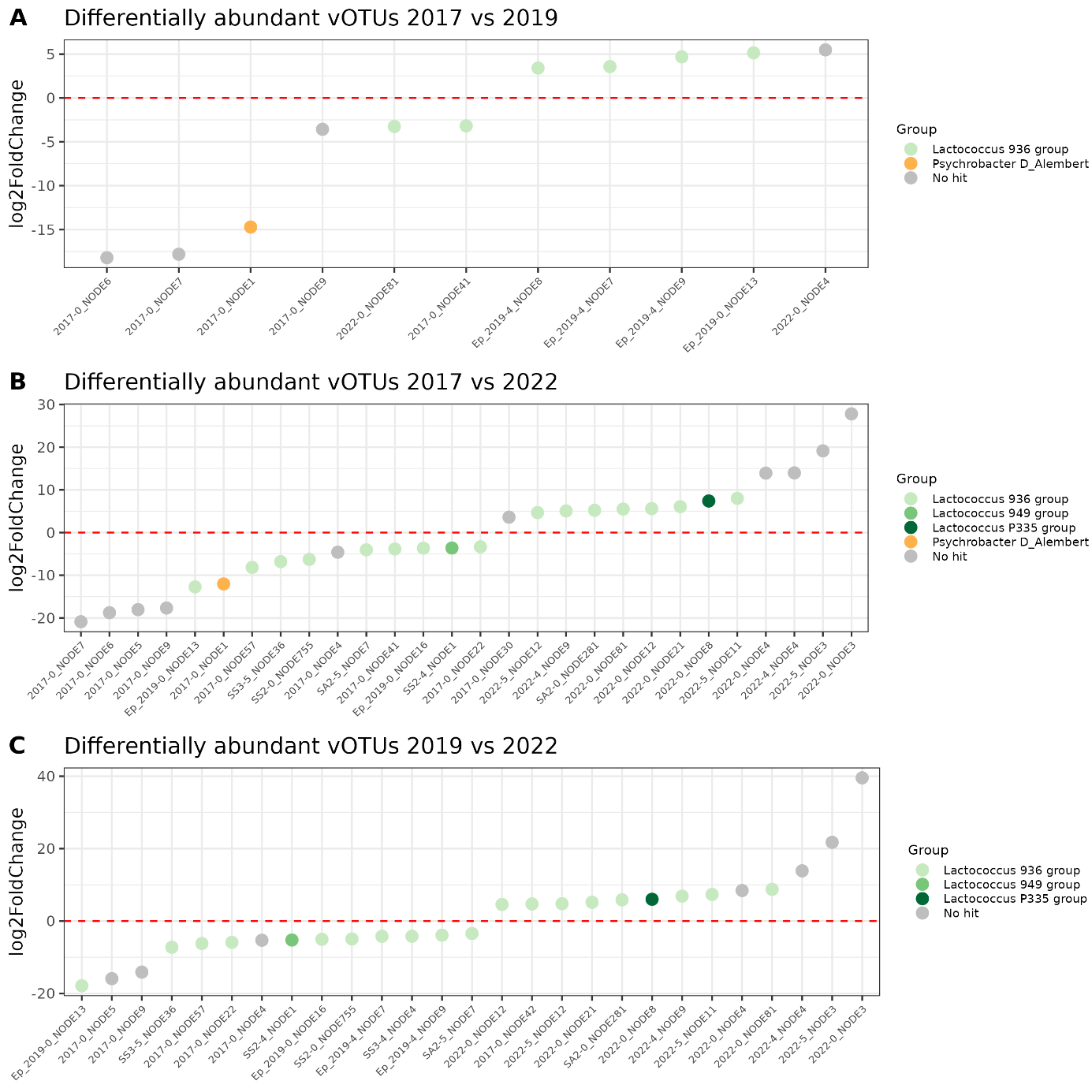

**Figure S3: Differential abundance analysis of the virome data between the three years of production (long-term study).** A) Comparison between years 2017 and 2019. B) Comparison between years 2017 and 2022. C) Comparison between years 2019 and 2022. Log2 fold change values obtained for differential abundant vOTUs (dots) are represented and the dot colour indicate the predicted host genus.
